# Biomolecular Resolution as a Quantification of Structural Information Content

**DOI:** 10.1101/2023.10.17.562748

**Authors:** Colin D. Kinz-Thompson, Korak Kumar Ray

## Abstract

Resolution is a poorly defined, *ad hoc* concept in structural biology, and multiple, seemingly conflicting definitions exist. In part, this ambiguity is because the structural information describing a biomolecule manifests in different ways depending upon the length scale being considered, while typical approaches to resolution just consider a single length scale. To address this shortcoming, here we develop a definition of resolution as a measure of how effectively those different manifestations of biomolecular structural information are embedded in experimental data across different length scales. We show how biomolecular structures can be decomposed into a hierarchy of structural elements, and combinations of those structural elements yield representations of the biomolecule with different degrees of spatial detail. A probabilistic measure, which we call “hierarchical resolution,” can then be calculated to determine the statistically optimal representation of a biomolecule in an experiment. Hierarchical resolution is a local, detection-sensitive resolution measure capable of quantifying the biomolecular structural information content of an experiment. As a case study, we use the bacterial ribosome to demonstrate how hierarchical resolution calculations relate to the reported experimental resolution. Altogether, these advances comprise a technique-independent definition of resolution and clarify the term “atomic resolution” for high-resolution techniques.

**STATEMENT OF SIGNIFICANCE:** Experiments probing geometric structure are an invaluable tool in understanding biomolecular function. While such experiments are primarily focused on determining atomic positions, other types of less-detailed, structural information also exist—even when the atomic information is obscured. Here, we present a theoretical framework for connecting the different types of biomolecular structural information and explain how to measure them. By determining when the different types of information are present, researchers can assess the reliability of their experiments. More significantly, this framework provides a solid theoretical basis for understanding and interpreting biomolecular structure across all length scales, which is evermore necessary as novel experimental techniques and artificial intelligence methods further enable researchers to study ever larger biomolecules in their natural, cellular contexts.

## INTRODUCTION

All molecules are governed by free energy landscapes that are shaped by the physical laws dictating atomic behavior (1). Large biomolecular complexes, however, are phenomenologically distinct from other types of molecules, as they perform the unique and complex tasks that sustain life. This emergent behavior is possible because the free energy landscapes of these complexes have been shaped over eons by evolutionary pressures to carry out biological functions and perform thermodynamic work (2). Since this is not the case for all molecules, it is clear that biological function has appeared as an emergent phenomenon at the molecular, rather than atomic, length scale (3).

The molecular-length-scale details that define and enable the biological function of a biomolecular complex are traditionally organized along a discrete hierarchy of structural information: primary, secondary, tertiary, and quaternary structure (4). Structural biology experiments, such as X-ray crystallography and single-particle cryogenic electron microscopy (cryoEM), are routinely able to extract structural information from every tier of this hierarchy—typically by building the best atomistic model (5–7) that explains certain experimental data (8). While the emergent, supra-atomic information that ultimately leads to biological function is intimately connected to atomistic definitions of structure and dynamics, it is worth noting that non-atomistic experimental techniques can still be used to obtain biomolecular structural information. For example, small angle X-ray scattering interrogates the global shape of a molecule (9), coarse-grained simulations provide information on the motion of helices and domains (10, 11), and generalized order parameters in nuclear magnetic resonance spectroscopy can localize thermodynamic information to individual residues (12). Moreover, some biomolecules, such as intrinsically disordered proteins, are considered to function as conformational ensembles that are statistically ill-defined on the atomic scale (13). Thus, biomolecular structural information both exists, and can be extracted, on multiple length scales.

The type of biomolecular structural information that can be extracted from different length scales is determined by the concept of resolution. For example, if a single-particle cryoEM experiment produces a low-resolution cryoEM density map, then building a coarse-grained model to analyze that biomolecular complex may be more appropriate than building an atomistic model (8). Doing so avoids overfitting atomic-scale structural information that is not embedded in the low-resolution data, while still allowing the extraction of existing structural information about the positions of the residues or helices. Unfortunately, it is unclear when structural information from a particular length scale is present or not in the data, in part because resolution is an ambiguous term.

In the traditional definition from the field of imaging, resolution is a binary property defined by whether two imaged objects are distinguishable in an experimental image (14). This definition is difficult to apply to structural biology experiments (15). That difficultly leads to cryoEM experiments, for example, relying on the gold-standard Fourier shell correlation (FSC) approach to explain resolution, which provides a robust experimental self-consistency but no structural insight into the biomolecule being investigated (16). Similarly, while “atomic resolution” could be a useful point of common reference for comparisons on the supra-atomic scale, even this term is ambiguous and often misused in structural biology studies (17, 18). As the continued development of techniques such as cryogenic electron tomography extend the scale of biomolecular investigation even further from the atomic realm (19), a precise definition of biomolecular structural information across all length scales, and an understanding of how it connects to resolution, is ever more important for alleviating this uncertainty.

A unifying theory connecting biomolecular structural information at different length scales to the concept of resolution is essential for understanding the supra-atomic structure, dynamics, and function of biomolecules. Here, we address the lack of such a theory by developing an unambiguous, model-based definition of biomolecular structural information. To do so, the traditional hierarchical tiers of biomolecular structure are used to quantitatively guide the interpretation of structural information at different length scales. The novelty of this approach is that the hierarchy of structure enforces both the presence and absence of all possible forms of structural information, thereby providing a clear meaning of resolution for biomolecules: the resolution of a structural dataset is determined by whether the presence of particular biomolecular structural elements with a specific length scales can be statistically justified. We show that this new approach of “hierarchical resolution” is functionally equivalent to the types of resolution criterion traditionally employed in imaging fields (14), but also accounts for experimental details, such as detection sensitivity and local data quality, by quantifying the biomolecular structural information content of the data. To demonstrate the utility of hierarchical resolution calculations, and their correspondence to our intuition about the term “resolution” as used in the fields of structural biology and biophysics, we used hierarchical resolution to analyze all the cryoEM-derived structures of the bacterial ribosome deposited in the Protein Data Bank (PDB). This analysis demonstrates the effectiveness of hierarchical resolution as a quantitative definition of atomic resolution, and, more generally, as a measure of the structural information content of structural biology experiments.

## MATERIALS AND METHODS

### Two-dimensional resolution measure simulations

The simulation and analysis of two-dimensional images were performed in Python; the corresponding code is available in a Zenodo repository (20). Two particles at a distance of 1 arbitrary unit (a.u.) apart were rendered as pixelated normal distributions onto a 13 x 13 pixel square grid with pixel dimensions 0.5 x 0.5 a.u.^2^. White noise was added to each pixel to achieve the desired peak signal-to-noise ratio (PSNR) for the simulation. Hierarchical resolution calculations were performed using *equal a priori* model priors, and *σ*_0,*min*_ = *σ*_1,*min*_ = 0.1 a.u., *σ*_0,*max*_ = 1.0 a.u., and *σ*_1,*max*_ = 10.0 a.u. priors for the marginalization of *σ_d_*, which was performed using Gaussian quadrature. 1000 images per imaging condition were simulated, and the median value of *P* under those conditions was used as an estimate for ⟨*P*⟩.

### Hierarchical resolution calculations on bacterial ribosomes

High-resolution bacterial ribosome structures were identified in the PDB (21) by searching for all entries: (1) with an experimental method of “ELECTRON MICROSCOPY;” (2) that contained the keyword “ribosome;” (3) that had a reported resolution less than 5 Å; (4) with a biological sample from a bacterial source organism; (5) where the entire complex was larger than 400 kDa; and (6) which had a corresponding deposition in the Electron Microscopy Data Bank (EMDB) (22). These search criteria yielded 925 entries released prior to October 1, 2026.

Simplified, isotropic hierarchical resolution calculations were performed with Python and C code that is available in a Zenodo repository (20). This implementation took approximately 40 ms per residue on a laptop computer. Polymer chains, excluding water, ions, and small molecules, were loaded from mmCIF files deposited in the PDB. CryoEM densities were loaded from MRC files deposited in the EMDB. Amino acid or nucleic acid residues of interest were identified as those containing a CA or C4’ atom name, respectively. For each residue of interest, a cubic (approximately 16 x16 x 16 Å^3^) volume of the quantized cryoEM density grid, centered around the uniformly weighted center of mass (COM) of the atoms in the residue, was extracted as a local *ε* -neighborhood for the calculation. Atomic-level *σ*_0_ dependent models, or residue-level *σ*_1_ dependent models of the residue were rendered on the grid of this local *ε*-neighborhood. In the former case, Gaussians with width *σ*_0_ centered at the location of each atom were integrated over each voxel within 5 *σ*_0_ of the location of each atom and summed together with unit weight. In the latter case, a single Gaussian with width *σ*_1_ centered at the COM of all the atoms was integrated over each voxel within 5 *σ*_1_ of the COM, and all voxels were weighted by the sum of the unit atom weights. At 101 log-spaced values of *σ_d_* between *σ_d_*_,*min*_ and *σ_d_*_,*max*_, the log-evidence value for each model calculated. Simpsons’ rule was used with these values to marginalize out the value of *σ_d_* from the joint probability of the evidence and the log-uniform prior for *σ_d_*. The resulting marginal-evidence values for the atomic-level and residue-level models were used as the model likelihoods in Bayes’ rule with equal probability model priors of 1/2 to perform the model selection calculation and calculate *P_m_* as the posterior value for the residue under consideration. These values, along with the log-evidences and values of *σ_d_* with maximum evidence for each residue exported into text files, which are available in the Zenodo repository (20).

## RESULTS

### The hierarchical decomposition of biomolecular structure categorizes structural information

The Richardson approach to schematic visualizations of biomolecular structure uses multiple graphical representations to simultaneously highlight the scope of structural information within a biomolecular complex (23, 24). This multi-resolution approach highlights the fact that unique structural features exist on different length scales. Inspired by this perspective, we hypothesized that the features used in Richardson-style schematics are quantifiable sources of emergent information concerning the structural organization of a biomolecular complex. To characterize that information, here we have developed a precise definition of a biomolecular structure as a compound, multi-resolution entity that simultaneously exists on several well-defined spatial length scales.

In this approach, the structure of a biomolecule is organized along a defined hierarchy of spatial length scales by making successively more granular approximations of its overall geometry (Fig. 1). Doing so yields a tiered set of structural elements that can be combined to form a hierarchical structural representation (HSR) of the biomolecule—each HSR representing a unique degree of detail. Such multi-resolution approximations are often used for efficient geometric computations in computer science, and can be understood as space partitioning trees (25). In general, such partition trees hierarchically decompose a region of high-dimensional space, creating a nested set of successively more sub-divided regions. The resulting decompositions act as progressively more resolved approximations of the original space. Each sub-graph of the partition tree (*i.e.*, an HSR) is then a unique approximation of the entire region (*i.e.*, the biomolecular complex) at a different level of geometric detail.

**Figure 1.**
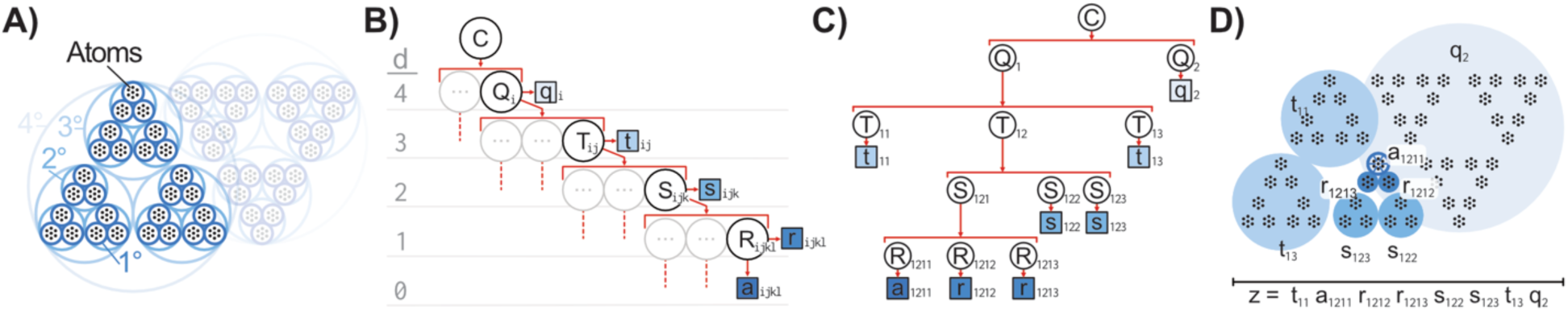
Hierarchical biomolecular approximations. (A) Schematic of bounding surfaces decomposing a hypothetical homodimeric complex with C_2_ symmetry into structural elements. Atom locations are shown in black. (B) A space partitioning tree to generate HSRs. Only one generic path is shown for clarity. Hierarchical level index, *d*, is shown to guide the eye. (C) Example HSR shown as a sub-graph of a partition tree. (D) Schematic diagram of the HSR, *z*, shown in panel c. Atom locations are present for clarity, but are not part of the HSR.

The process of creating a partition tree for a biomolecular complex begins by constructing a bounding surface around the entire complex, and then adding sub-dividing boundary surfaces to separate each constitutive molecule within the complex (*i.e.*, at the quaternary structure level) (Fig. 1A). Each of the molecules in the complex can be further decomposed into domains by adding more sub-dividing boundary surfaces (*i.e.*, at the tertiary structure level). The entire biomolecular complex can then be thought of as either a collection of molecules, a collection of domains, or a mixed collection of molecules and domains—each collection is an HSR of the entire complex. Likewise, the domains can be further decomposed into helices and loops (*i.e.*, at the secondary structure level), which can be further decomposed into residues (*i.e.*, at the primary structure level) that ultimately can be decomposed into atoms (Fig. 1A). The successive, hierarchical decompositions of each structural element in the biomolecular complex are mapped by the partition tree (Fig. 1B). HSRs are then created by traversing down the partition tree to select a complete collection of potentially mixed length-scale structural elements to define the approximation geometry of the complex (Fig. 1C-D). The set of all possible HSRs defines the scope of all possible geometric approximations and thus contains all the structural information about that biomolecular complex.

Once a partition tree is constructed, an HSR can more readily be generated by transforming the partition tree into a probabilistic context free grammar (PCFG) (26). The PCFG provides a set of rules that define when, and how often, the different structural elements should appear in an HSR, ensuring that it obeys the hierarchy imposed by the partition tree. In this work, the structure of a biomolecular complex, *C*, is considered on five distinct, hierarchical length scales: quaternary (*Q*), tertiary (*T*), secondary (*S*), residue (*R*), or atomic (*A*); these length scales are indexed with *d* = 4, 3, 2, 1, and 0, respectively. For a partition tree based on this hierarchy of length scales, the equivalent PCFG rules are:

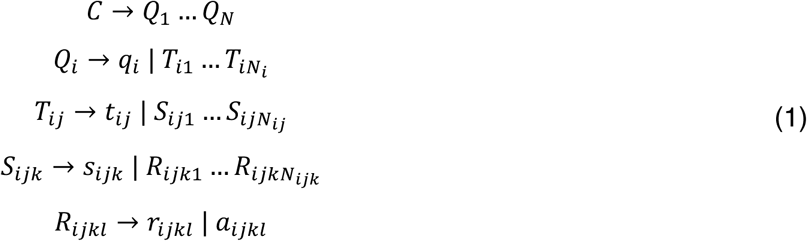

where the upper-case letters are non-terminal symbols that can be further subdivided, the lower-case letters are terminal symbols that cannot be further subdivided, and the various *N* denote the number of child structural elements created when a parental structural element is subdivided. Each rule, ℛ, in the PCFG (Eqn. 1) captures how a structural element is subdivided within the partition tree. Traversing a partition tree to create an HSR can equivalently be thought of as building a sentence of terminal structural elements picked using the rules of the PCFG. The probability of each ℛ being enacted has an associated accessibility, *ψ*_ℛ_ = *e*^−*βE*ℛ^ , where *β* is the coolness and *E*_ℛ_ is the potential of enacting that rule. A complete HSR is thus created by beginning with the biomolecular complex, *C*, picking a rule consistent with the *ψ*_ℛ_, and continuing until all non-terminal symbols have been replaced, yielding the probability of this HSR, *z*, as *P*(*z*) ∝ ∏_ℛ∈*z*_ *ψ*_ℛ_.

The resulting HSR is an ordered sentence of structural elements that represents the entire biomolecular complex, *z =* (*ξ*_1_, …, *ξ*_|*z*|_). For example, the HSR *z =* (*t_11_, a_1211_, r_1212_, r_1213_, s_122_, s_123_, t_13_, q_2_*) represents a small homodimeric complex where each protomer has three domains (Fig. 1C-D). These domains are not resolved in the second protomer (*q_2_*). In the first protomer, two of the domains are resolved at the tertiary level (*t_11_* and *t_13_*), while the remaining domain is further resolved to “lower” levels (*d < 3)*. Specifically, this domain is resolved into two secondary structure elements (*s_122_* and *s_123_*). The residual element, which would otherwise be represented by *s*_121_, is further resolved down to the atomic level for one residue (*a_1211_*), but only to the residue level for *r_1212_* and *r_1213_* (Fig. 1C-D).

Conceptually, each HSR is a statement of a researcher’s structural understanding of a particular biomolecule. The HSR describes which structural information is relevant in an experiment. As such, HSRs are not coarse-grained models, *per se*. Instead, they exist in an information space, and a forward model is required to transform them into structural elements in real space. This difference between an HSR and a coarse-grained model is more evident when considering the information statistics of the PCGF (26), which are different than the statistical mechanics of a coarse-grained molecule (27, 28). The probability of a particular HSR depends on the *E*_ℛ_ potentials of the PCFG, not the kinetic and potential energy of the atoms in the molecule. For the hierarchy described above, the total number of distinct HSRs is 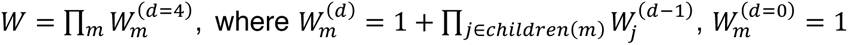, and *m* indexes the structural elements at level *d*. For a prototypical 300-residue-long protein that is decomposed into three domains, each with 10 secondary structure elements that each contain 10 residues, then there are approximately *W* ≈ 2^1⋅3⋅10⋅10^ = 2^300^ ≈ 10^90^different HSRs. Fortunately, the inside-outside algorithm can be used for marginalizations and probabilistic inference—in particular to determine the *ψ*_ℛ_ and *E*_ℛ_ for the different rules of the PCFG, as done in linguistics (29). Furthermore, once inferred, it is possible to treat the PCFG as a hierarchical hidden Markov model to generate high-probability HSRs (30).

In practice, modern structural biology techniques simplify the information statistics of these PCFGs. For example, the resolution revolution in single-particle cryoEM (31) has eliminated many low-resolution HSRs from having an appreciable probability for modern cryoEM experiments. Specifically, the potentials for all the PCFG rules that yield terminals above the residue level (*i.e.*, *d* > 1) are effectively infinite in a successful experiment, such that the corresponding marginal probabilities are *P*(*Q_i_* → *q_i_*) = *P*F*T_i_*_j_ → *t_i_*_j_G = *P*F*S_i_*_j*k*_ → *s_i_*_j*k*_G = 0 . In this case, the only HSRs worth considering are those defined exclusively at the residue or atomic level (*i.e.*, *d ≤ 1*). There are exactly *W* = 2*^N^* of such HSRs, where *N* is the number of residues in the biomolecular complex. From the inside-outside algorithm (29), each branch of this reduced partition tree is then independent, and so the structural elements representing each residue are independent of the others for these HSRs. Thus, *P*(*R_m_* → *a_m_*) = (1 + *e*^−*β*Δ*E*^*^m^*) ^-1^ *e*^−*β*Δ*E*^*^m^*, where Δ*E_m_* is the potential difference for the *m^th^* residue being resolved at the atomic-level (*a_m_*) *versus* at the residue-level (*r_m_*). By assuming the residues are distinguishable, independent sites, the partition function for the PCFG is *Z* = ∏*_m_*(1 + *e*^−*β*Δ*E*^*^m^*) ^-1^, and thus the information entropy of the complete set of HSRs is

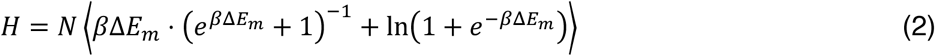

where the average is over all *N* residues.

Altogether, the information statistics of these hierarchical approximations of biomolecular structure suggests that structural biology experiments function, at least in part, by providing data that allows a researcher to determine which HSRs are the highest probability representations of the biomolecular structure. The analysis of that data consequently yields a new PCFG with altered *E*_ℛ_ potentials; we refer to such a PCFG from the same partition tree but with different *E*_ℛ_ potentials, as a structural hierarchy, ℋ. Each ℋ has an information entropy (Eqn. 2), which quantitatively represents the researcher’s uncertainty about the structural details of the biomolecule being investigated in the experiment. When hot enough (*β*Δ*E_m_* → 0), Eqn. 2 reduces to the expected *H* = *N* ln 2 in which the potentials are completely washed out by thermal energy. This maximum-entropy situation represents the case where a researcher has no knowledge about the biomolecular structural information of the complex being studied other than what is provided by the laws of physics used in defining the partition tree (32). Moreover, this maximum-entropy case provides an appropriate structural hierarchy for the initial situation prior to performing the experiment, ℋ*_i_*. After an experiment, the experimental data informs upon updated *E*_ℛ_ potentials to yield a final structural hierarchy, ℋ*_f_*. The transition from ℋ*_i_* to ℋ*_f_* following a completed experiment results in a loss of information entropy, Δ*H* = *H_f_* − *H_i_* ≤ 0, or rather an increased certainty about which structural details most accurately represent the biomolecule in the experiment. Thus, as a comparison of the structural information in ℋ*_i_* and ℋ*_f_*, Δ*H* quantifies the structural information content of the experiment.

#### Inferring the best hierarchical structural representation

Probabilistic inference (33), which is a particularly effective approach for analyzing structural biology experiments (34–37), can be used to statistically determine which HSR is the best representation of an experiment. This calculation requires a forward model to project HSRs into the experimental space where they can be probabilistically compared on the basis of how well each explains the data—a process we call a “hierarchical resolution” calculation. As a general method, hierarchical resolution is applicable to imaging-style structural biology experiments, such as single-particle cryoEM, super-resolution fluorescence microscopy, or atomic-force microscopy. Here, we explain the general approach using an isotropic forward model to quantify the structural information content of an experiment.

To quantitatively make a comparison between different HSRs, the probability that a particular HSR, *z*^C^ ∈ {*z*}, where {*z*} is the set of all HSRs for a particular biomolecular complex, represents the biomolecular complex in some experimental data, *D*, can be calculated using Bayes’ rule as

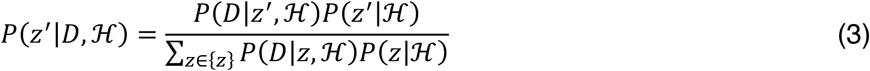

Here, *P*(*x*|*y*) is the conditional probability of *x* given the value of *y*, and the conditional nature of these probabilities on the specific structural hierarchy ℋ represents the partition tree and *E*_ℛ_ potentials of the PCFG (Fig. 1). When ℋ specifies *equal a priori* model prior probabilities, *P*(*z*|ℋ), this calculation becomes a comparison between the marginal likelihoods, *P*(*D*|*z*, ℋ), for all of the HSRs defined by the partition tree.

Calculating any of those marginal likelihoods requires a forward model to project the HSR into the dataspace of the experiment. Forward models are inherently subjective and depend upon the desired level of physical complexity. Here, we use a simple forward model where linear imaging of atoms yields isotropic Gaussians—this is a reasonable approximation for techniques such as single-particle cryoEM (38–40). Structural elements from higher levels of the hierarchy (*d* > 0) are then imaged as isotropic super-atoms centered at the COM of the constitutive atoms. As a concept, hierarchical resolution does not depend upon the specific choice of forward model. For instance, an anisotropic forward model would be more physically realistic; however, it is currently unclear how to easily generalize the anisotropic shapes encountered at the secondary structure levels and higher (*d* > 1).

The data in an imaging-based structural biology experiment, such as the cryoEM density map created during a single-particle reconstruction, is a noisy set of discrete densities {*Y*(*v⃗*)|*v⃗* ∈ *G*}, where *G* is a spatial grid and *v⃗* is a grid voxel position. Applying the isotropic forward model to an HSR, *z*, evaluated on *G* by integrating over the voxel *v⃗*, yields a discretized spatial density

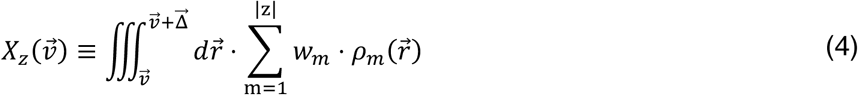

where ^g^Δ⃗ is the dimensions of the voxel, and *r⃗* is the position in real space. In this simple case, the density kernel for the *m^th^* structural element is *ρ_m_*(*r⃗*) = N(*r⃗*|μ⃗*_m_*, *σ*^2^_m_), which is an isotropic, multivariate normal distribution (N) centered at μ⃗*_m_* with variance *σ*^2^ , where *m* indexes all the structural elements in *z* = F*ξ*_1_, … , *ξ_m_*, … , *ξ*_|*z*|_G. The weight of each density kernel is taken as the imaged contribution from all constitutive atoms in that structural element, *w_m_* = ∑*_i_*_∈£_*_m_ω_i_*, where *ω_i_* is the individual contribution for the *i^th^* atom. For example, in single-particle cryoEM, *ω_i_* defines how the atomic scattering factor for the *i^th^* atom appears in the Coulombic interaction potential with an incident electron plane wave in the microscope (39–42).

Eqn. 4 yields a set of unscaled, unitless densities {*X*(*v⃗*)|*v⃗* ∈ *G*}, which we call a “shape template” because it captures the latent structure of the biomolecule encoded as correlations of the density between the voxels (43). Assuming the imaging process induces unknown distortions of positive scale (*m*), offset (*b*), and additive white noise (*τ*) to the shape template, the experimental data is distributed as *Y*(*v⃗*) ∼ *mX*(*v⃗*) + *b* + N(0, *τ*^−1^). From the point-of-view of understanding biomolecular structure, *m*, *b*, and *τ* are nuisance parameters—they do not convey useful molecular information. These nuisance parameters can be analytically removed by marginalization (43), yielding the marginal likelihood of observing the dataset, regardless of any distortions by scale, offset, or noise,

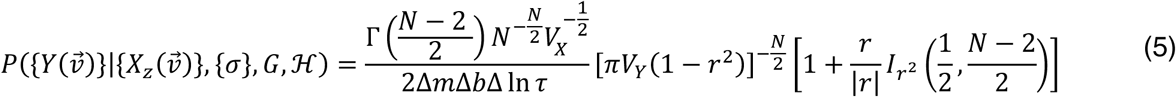

where Γ(*a*) is the gamma function, *I_v_*(*α*, *β*) is the regularized incomplete beta function, *N* is the number of voxels in *G*, the variances of {*X*(*v⃗*)} and {*Y*(*v⃗*)} are *V_x_* = ⟨*X*^2^(*v⃗*)⟩ − ⟨*X*(*v⃗*)⟩^2^ and *V*_F_ = ⟨*Y*^2^(*v⃗*)⟩ − ⟨*Y*(*v⃗*)⟩^2^, respectively, and *r* = (⟨*X*(*v⃗*)*Y*(*v⃗*)⟩ − ⟨*X*(*v⃗*)⟩⟨*Y*(*v⃗*)⟩)/}*V_X_V*_F_ is the correlation coefficient between {*X*(*v⃗*)} and { *Y*(*v⃗*)}. The Δ*m*, Δ*b* , and Δ ln *τ* are defined as Δ*f*(*x*) ≡ *f*(*x_max_*) − *f*(*x_min_*), and arise from the independent uniform or log-uniform prior probability distributions used for the nuisance parameters (43). These prior probability normalization factors do not need to be precisely defined, because they cancel in Eqn. 3 during a hierarchical resolution calculation.

To calculate the full marginal likelihood *P*(*D*|*z*, ℋ) used in Eqn. 3, the conditional dependence on the widths of the isotropic density kernels, {*σ*}, has to be removed from Eqn. 5. This is achieved by marginalization of the joint probability distribution *P*({*Y*(*v⃗*)}, {*σ*}|{*X_z_*(*v⃗*)}, *G*, ℋ) = *P*({*Y*(*v⃗*)}|{*X_z_*(*v⃗*)}, {*σ*}, *G*, ℋ) ⋅ *P*({*σ*}|ℋ). A facilitating assumption we apply in hierarchical resolution is that every structural element from the same tier of the structural hierarchy has the same value of *σ*, such that *σ_m_* = *σ_d_* for every *ξ_m_* on the *d^th^* tier of the partition tree (Fig. 1b). Assuming that *σ_d_* are independent and each is distributed according to the maximum entropy distribution for an unknown magnitude, the prior probability distribution for {*σ*} is

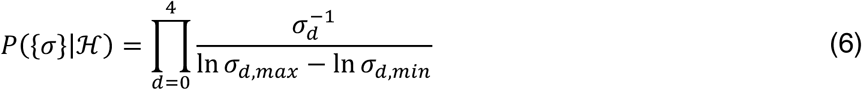

where *σ_d_*_,*min*_ and *σ_d_*_,*max*_ are the minimum and maximum values of *σ_d_* supported by the prior probability distribution (see below). The resulting marginalization integral is

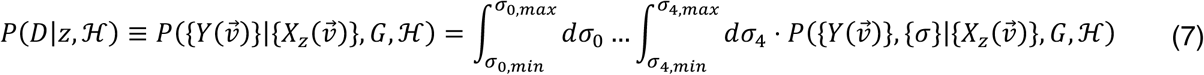

For computational tractability, these integrals can be estimated on a grid as a Riemann-style sum. Following marginalization of each HSR using Eqn. 7, Eqn. 3 may then be used to determine the probability that an HSR is the best structural description of the biomolecular complex imaged in the experiment. In comparison to the model prior probabilities, *P*(*z*|ℋ) , the resulting model posterior probabilities, *P*(*z*|*D*, ℋ), serve as quantitative measurements of the structural information acquired during an experiment. For example, if the HSR where all the *ξ_m_* = *a_m_* has *P*(*z*|*D*, ℋ) = 1, then the experiment unambiguously yielded atomic information for every residue in the biomolecular complex. Alternatively, if the probability of every HSR in {*z*} is the same, then the experiment was completely ambiguous and yielded no structural information.

Finally, it is again worth highlighting that the resultant probabilities are conditional upon the choice of structural hierarchy, ℋ. As such, the choice of partition tree is particularly fundamental to the results of these calculations. In this sense, the researcher’s knowledge of the composition (*e.g.*, primary sequence, subunit stoichiometry) and hierarchy of structure (*e.g.*, secondary structure assignments, domain decompositions) of the biomolecular complex is integral to any conclusions concerning structural elements. Fortunately, nearly all biomolecular studies require such information to some degree, and the field has developed many experimental and computational methods of making such determinations. For example, domain decompositions are often performed by sequence homology-based classification (*e.g.*, using profile hidden Markov models (44)) in protein sequence databases, as is done with InterPro (45). Secondary structure assignments have been performed for decades using classic methods, such as DSSP (46) or STRIDE (47) for proteins, or Mfold (48) for nucleic acids. Notably, recently developed machine learning-based protein language models can predict both the secondary structure assignments and tertiary structure from the primary sequence alone, as is done by ESM3 (49). Thus, in addition to literature consensus concerning structural elements for a biomolecule of interest, a researcher has many options to help develop the partition tree to use in hierarchical resolution. An exciting prospect for future development of this framework, however, is to compare the results from different hierarchies—something that is essential for assessing the role of conformational dynamics in defining resolution (see *Discussion*).

### Hierarchical structure and the bounds of resolution

External factors, such as microscope hardware or molecular sample heterogeneity, affect whether a particular structural element within an HSR will be useful for describing the biomolecular structural information in an experiment. These factors are related to our understanding of “resolution,” because they affect whether a structural element will appear in the data, or whether it will appear to have been replaced by its higher-level parent element. For example, if two atoms are spatially close together and the nominal resolution of a cryoEM density map is too poor, then the atoms can *appear* to merge into a single entity in the map. In such a case, the HSR containing a residue-level representation of these atoms may be more appropriate than one with atomic-level representations.

In imaging fields, the Rayleigh resolution criterion describes the two-dimensional equivalent of this phenomenon. It defines a cutoff distance where two point-objects *appear* to merge into one. At distances shorter than the cutoff, the point-spread function (PSF) of the imaging system induces such a significant apparent overlap in an image that the objects are not considered distinguishable from one another; at distances longer than the cutoff, the objects are considered to be resolved (14). In this sense, there is a deep connection between the role that cutoff distances play in defining the concept of resolution, and the determination of when a lower-level structural element elements is no longer a physically coherent, nor appropriate, description of biomolecular structural information in hierarchical resolution.

The connection between resolution and the structural hierarchy is that each tier of the structural hierarchy (Fig. 1B) naturally encodes a “Rayleigh-like” resolution criterion for its structural elements. These Rayleigh-like resolution criteria naturally emerge from the prior probability distribution, *P*({*σ*}|ℋ) (Eqn. 6). Setting the *σ_d_*_,*max*_ is equivalent to determining how large a particular structural element can appear before it loses physical meaning and a structural element from its parental tier (*i.e.*, d+1) becomes the *de facto* appropriate representation for features of that size. In this work, *σ_d_*_,*max*_ is defined as the value of *σ_d_* when the shell encompassing 50% of the isotropic Gaussian density of one structural element at level *d* intersects the center of another located a reference distance, *l_d_*, away (Fig. 2). This definition accounts for the three-dimensional nature of the structural elements. Consequently, hierarchical resolution enforces a three-dimensional Rayleigh-like resolution criterion in which structural elements are only considered resolved when the 50%-density shell for one object will not go past another reference element (Fig. 2B). This definition is motivated by the relative value of the three-dimensional density at the midpoint between two objects (0.744), which is very close to the generalized Rayleigh criterion cutoff value (0.735) for point-objects in two-dimensional optical microscopy (14).

**Figure 2.**
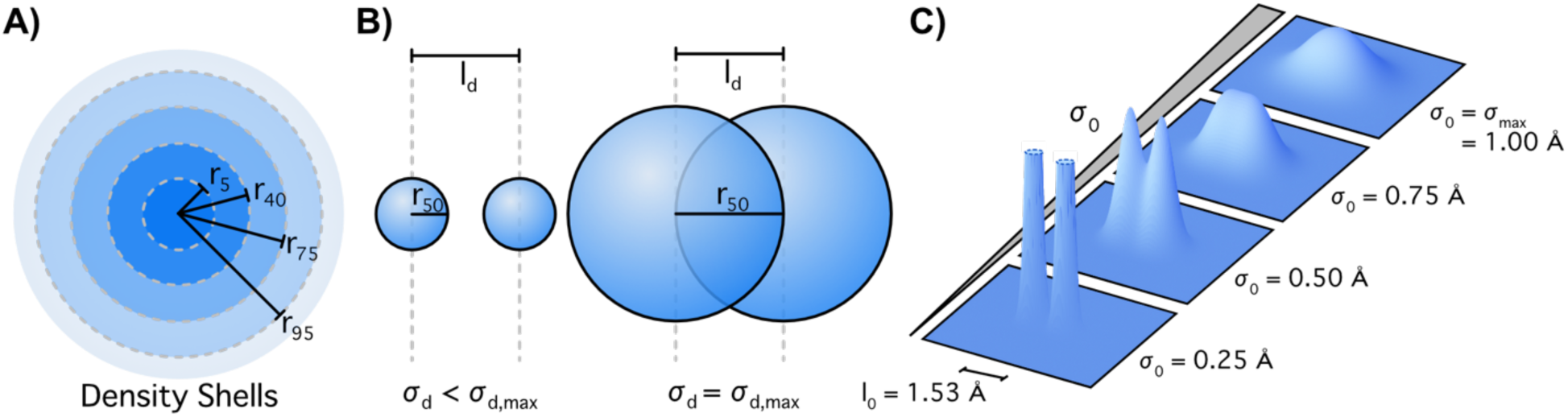
Hierarchical level-dependent resolution criteria in 3D. (A) Schematic cross section of a density kernel *ψ_m_* with radial density shells *r_x_* denoting the distance at which *x*% of the density is encapsulated. (B). Schematic of neighboring structural elements represented by narrow (left) and broad (right) density kernels. Significant overlap occurs when *σ_d_* causes *r*_50_ to equal the separation distance, *l_d_*. (C) Demonstration of two carbon atoms that are part of a C-C bond imaged with increasing *σ*_0_ until they appear to merge at *σ*_0_ = 0.995 Å.

Prior chemical knowledge must be used to define the reference distance, *l_d_*, that is used to set the corresponding *σ_d_*_,*max*_ . For the lowest level, *d = 0*, where the structural elements representing a residue are the individual atoms, the carbon-carbon single (C-C) bond is a reasonable reference distance. The C-C bond length in saturated hydrocarbons is *l*_0_ = 1.53 Å (50). For two isotropic Gaussian-distributed densities separated by 1.53 Å, the 50% overlap point occurs when the corresponding width *σ*_0_ = 0.995 Å. Thus, the choice of the C-C bond as *l*_0_ naturally leads to the determination that *σ*_0,*max*_ = 1 Å for all atomic profiles at the *d = 0* level of the structural hierarchy (Fig. 2C).

For the *d = 1* level, there are many reasonable reference distances that could be used to define *σ*_1,*max*_. For instance, the typical distance between nucleic-acid residues stacked along a B-form helix is 3.4 Å, the typical C_ɑ_-C_ɑ_ distance between consecutive amino-acid residues in a protein is 3.8 Å (51), and the typical distances between the two strands in a β-sheet or ɑ-helix are 4.7 Å and 5.4 Å, respectively (4). An empirical solution is to use the average distance to the closest neighboring residue as calculated across all high-resolution biomolecular structures in the PDB. For high-resolution structures, the average COM to nearest-neighbor COM distance is 4.33 Å (see *Supporting Material;* Fig. S1). For two isotropic Gaussian-distributed densities separated by 4.33 Å, the 50% overlap point occurs when *σ*_1_ = 2.82 Å. Thus, the empirical choice of *l*_1_ = 4.33 Å naturally leads to the determination that *σ*_1,*max*_ = 2.82 Å for all residue profiles at the *d = 1* level of the structural hierarchy.

The ease with which the *d = 0* and *d = 1* cutoff widths were defined comes from the fact both atoms and residues are reasonably isotropic in comparison to higher-level structural elements, such as helices. Therefore, higher levels of the hierarchy require more complex, anisotropic resolution criteria be further developed. For high resolution experiments, it is unlikely that the experimental data will require an

HSR with higher-level structural elements (*i.e.*, *d > 1*). In that case, the exact specification of those *σ_d_*_,*max*_ is unnecessary, because model priors with *P*(*z*|ℋ) = 0 for every *z* that contains a *ξ_m_* with d > 1 should be employed anyway. This is equivalent to using a PCFG with *P*(*Q_i_* → *q_i_*) = *P*F*T_i_*_j_ → *t_i_*_j_G = *P*F*S_i_*_j*k*_ → *s_i_*_j*k*_G = 0, as discussed above.

The Rayleigh-like resolution criteria only govern the upper limits that define the structural elements (*σ_d_*_,*max*_). Consequently, without invoking other physical principles (32), the lower limits are agnostic to the effects of the hierarchy and thus all *σ_d_*_,*min*_ should be the same. The smallest possible value of *σ_d_*_,*min*_ occurs when broadening is caused only by thermal motion (*e.g.*, with no reconstruction errors or conformational heterogeneity). An estimate of *σ_d_*_,*min*_ can be made from a best-case scenario by noting that a well-behaved, plunge-frozen crystal of metmyoglobin at 80 K had an overall B-factor of ∼5 Å^2^ (52), which corresponds to 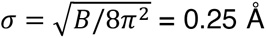. Given such an ideal sample and experimental conditions, it is unlikely to ever observe any structural element appear much smaller than this size in an experiment. Consequently, an order-of-magnitude estimate of *σ_d_*_,*min*_ = 0.1 Å is reasonable choice for every hierarchical level.

Conditioned upon a reasonably high nominal resolution structural experiment, such as in modern single-particle cryoEM (31), each residue is independent in every HSR (see above). Consequently, the marginal likelihoods factorize as *P*(*D*|*z*, ℋ) = ∏*_m_ P*(*D*|*ξ_m_*, ℋ). This factorization enables Eqn. 3 to be used to efficiently calculate the probability that the *m^th^* residue has only atomic resolution or residual-level resolution. Since *ξ_m_* = *a_m_* or *ξ_m_* = *r_m_* are mutually exclusive options for an HSR, the local probability of atomic resolution, *P_m_*, for the *m^th^* residue is derived from Eqn. 3 as

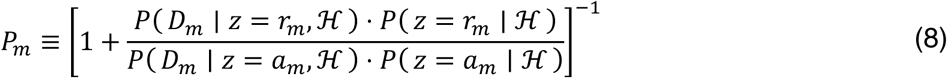

where *D_m_* is a local *ε*-neighborhood of the experimental data *D* (*e.g.*, within a cube with side length 2*ε* that is centered at the COM of the *m^th^* residue). Such localized calculations work by assuming that data outside the *ε*-neighborhood are distributed according to some (unspecified) model that is independent from the data inside the *ε*-neighborhood, so that those contributions cancel in Eqn. 3 (see Ref. (43) for more details). Thus, Eqn. 8 allows for a calculation of a local probability that a residue has atomic information, which is useful for experiments that exhibit, for example, preferred orientation complications.

#### Hierarchical resolution is a detection-sensitive resolution measure

To understand how hierarchical resolution quantifies biomolecular structural information, the performance of hierarchical resolution was compared to traditional resolution measures. Traditionally, both the Rayleigh resolution criterion and the Abbe limit are used to assess resolution in two-dimensional imaging experiments. They both specify a minimum distance between two objects, beyond which the two particles are considered resolved and can thus be separately observed in an image (14). Specifically, the Rayleigh resolution criterion determines when two point-particles are close enough that they appear to overlap in the resulting image, while the Abbe limit specifies when an aperture in the back-focal plane creates a hard limit to which spatial frequencies can be transferred into an image. The cutoff for the Rayleigh approach occurs when the separation between the two point-particles is *l*_R*aylei*gℎ_ = 0.61 *λ*/*NA* when imaged using a lens with numerical aperture *NA* and light of wavelength *λ*. The cutoff for the Abbe approach occurs when the separation is *l_Abbe_* = 0.5 *λ*/*NA* . For reference, *σ_xy_* = 0.21 *λ*/*NA* is the effective Gaussian width of a diffraction limited spot (14, 53). In comparison, whereas the Rayleigh resolution criterion and the Abbe limit make hard determinations about whether two structural elements are considered resolved, hierarchical resolution calculations yield a soft determination. If the probability for an HSR with two structural elements is high enough, then those two structural elements can be considered as resolved within the data.

To compare these three approaches, images of a simplified two-dimensional biomolecule that is composed of one residue made up of two atoms separated by a distance, *l*, were simulated. The only possible HSRs for this rudimentary molecule are {*z*} = {(*r*_1_), (*a*_1_)}; all higher-level *d > 1* structural elements are redundant due to the small size. The atoms were represented by pixelated isotropic Gaussians of unit weight in the simulated images, and a set of *N* = 1000 images, {*D*}, was simulated under different imaging conditions. Varying amounts of white noise with strength *σ*^2^ was added to each image, such that the peak signal-to-noise ratio (PSNR) varied from 0.5 to 10, where *PSNR* ≡ *D_max_*/*σ_sim_*, and *D_max_* is the maximum pixel intensity in the image before adding noise (Fig. 3A-B). These simulations were performed as a function of the normalized PSF width, *λ*/(*l* ⋅ *NA*), where a smaller width leads to more resolved images of the atoms, and a larger width leads to more diffuse images of the atoms that overlap and ultimately become unresolved. Subsequently, Eqn. 8 was used to estimate the average probability that the atoms in the molecule were resolved at the atomic level, ⟨*P*⟩ ≡ ⟨*P*( *a*_1_ ∣ *D*, *H* )⟩, for each simulated imaging condition. Plotting ⟨*P*⟩ *versus* the normalized PSF width enabled a direct comparison of hierarchical resolution *versus* the Rayleigh resolution criterion and the Abbe limit (Fig. 3C).

**Figure 3.**
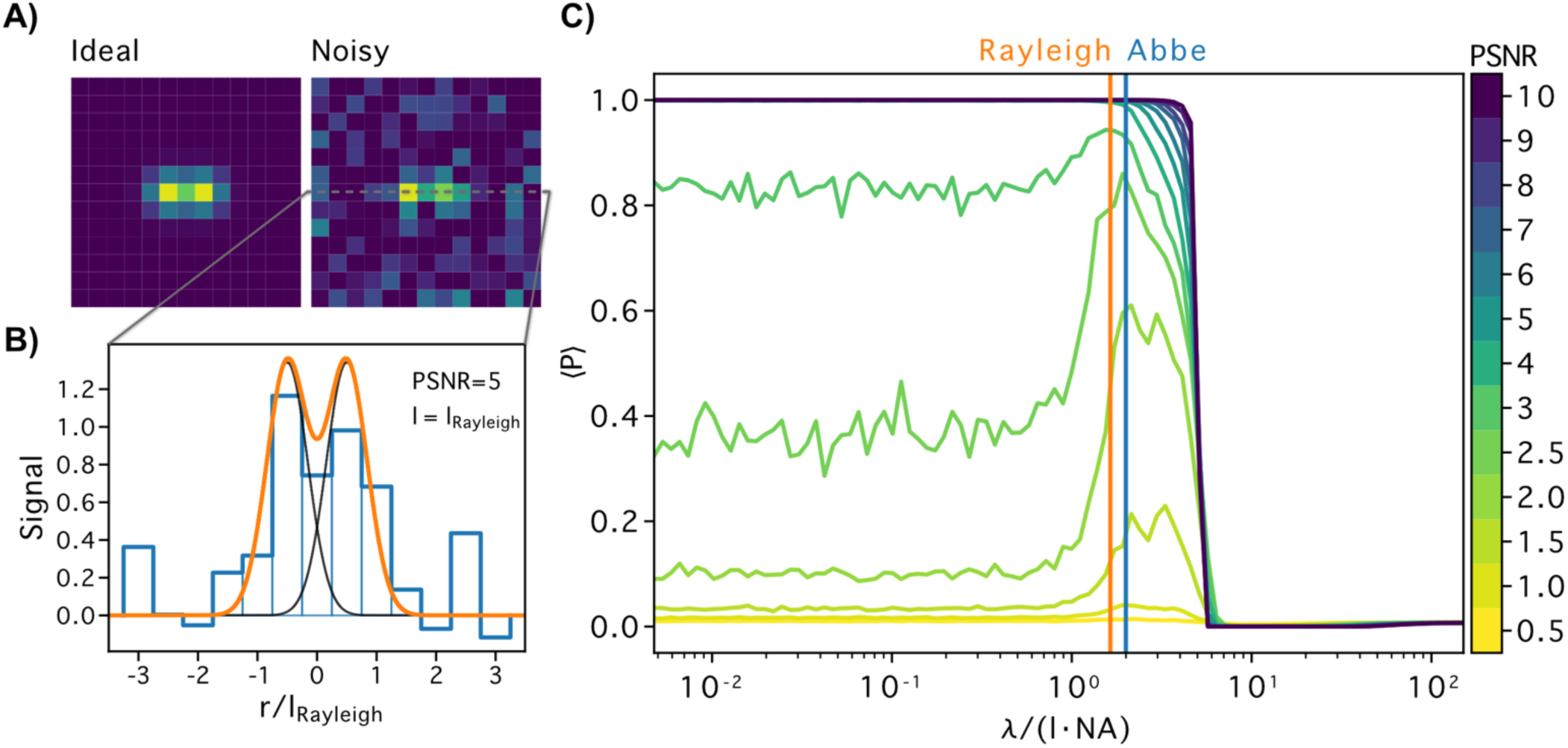
Comparison of hierarchical resolution to the Rayleigh resolution criteria and the Abbe limit. (A) Example simulated image of two atoms separated by a distance at Rayleigh limit with (right) and without (left) added noise. (B) Cross section of noisy, simulated image (blue) along inter-atomic axis. The non-pixelated ground truth is shown in orange, with individual contributions in black. (C) Plot of atomic resolution *vs.* imaging wavelength in simulated 2D images at various levels of noise. Vertical lines mark the Rayleigh (orange) and the Abbe (blue) limit of resolution.

Foremost, these simulations show that hierarchical resolution exhibits the same limiting behavior as the Rayleigh and Abbe approaches, which is expected of any resolution measure. ⟨*P*⟩ remains constant with increasing normalized PSF width until it sharply drops to zero at a normalized PSF width of ∼5. This drop defines a cutoff point after which the atoms are no longer resolved and the residue-level

HSR *z = (r_1_)* becomes the best representation of the biomolecule in the image. Notably, this drop-off occurs when *σ_xy_* ≈ *l*, where the separation between the atoms is the effective Gaussian width of a diffraction-limited spot. The equivalent Rayleigh and Abbe cutoff points occur earlier at slightly sharper normalized PSF widths of ∼1.6 and 2. Thus, the two-dimensional hierarchical resolution calculation provides slightly improved resolving power. The origin of this improved performance seems not to be the mismatch of the isotropic forward-model to the apparent anisotropic elongation along the inter-atomic axis as the atoms appear to merge. Instead, it seems to be related to the value of *σ*_0,*max*_, which was not fine-tuned for the simplified two-dimensional biomolecule situation here (see *Supporting Materials*; Fig. S2). However, we note that model-based resolution measures for two-dimensional fluorescence imaging have also found an approximately 3-fold increased performance over the Rayleigh criterion (54, 55).

While ⟨*P*⟩ is a resolution measure like the Rayleigh and Abbe approaches, it also accounts for detection sensitivity. Neither of the traditional resolution measures accounts for noise or detection sensitivity. Thus, while certain high frequency features might hypothetically be transferred into an image based on these cutoffs, the signal-to-noise ratio can be so poor that those features might not be observed despite them being “resolved.” Including detection sensitivity in a definition of resolution requires a model-based approach to describe the imaging and detection process (54, 55), and notable advances have been made for single-molecule fluorescence imaging (56, 57). For hierarchical resolution at small normalized PSF widths (*i.e.*, < 1), the value of ⟨*P*⟩ monotonically increases with increasing PSNR, demonstrating that that ⟨*P*⟩ directly relates to detection sensitivity (Fig. 3C). Since these curves saturate to ⟨*P*⟩ = 1 at PSNR of approximately 4, this calculation demonstrates that hierarchical resolution interprets low PSNR images as having only small amounts of statistical evidence for both the atomic- and the residue-level HSRs—neither HSR is conclusively supported. At a more reasonable PSNR (*e.g.*, see PSNR = 5 in Fig. 3A-B), ⟨*P*⟩ exhibits limiting behavior analogous to a binary resolution measure. Thus, hierarchical resolution is a detection-sensitive resolution measure, but one specifically for imaging-based experiments investigating three-dimensional biomolecular structure.

### Quantifying the atomic information content of the bacterial ribosome using hierarchical resolution

To establish the effectiveness of the biomolecular structural information quantification performed in a three-dimensional hierarchical resolution calculation, we analyzed all the “high-resolution” cryoEM-derived bacterial ribosome structures deposited in the PDB (21). Specifically, we sought to understand how atomic-scale information emerges as a function of the reported FSC-based resolution of single-particle cryoEM experiments. The choice of a single biological system reduced effects of molecular heterogeneity in complicating the interpretation. Moreover, as a large ribonucleoprotein complex containing both nucleic acids and proteins, the ribosome provided a chemically comprehensive scope for this investigation. Due to its high stability and large size (∼2.5 MDa), the bacterial ribosome has been the quintessential biomolecular system used for the development of single-particle cryoEM as a technique (58). Together with its biological significance, this has resulted in many single-particle cryoEM studies of the bacterial ribosome throughout the past few decades at all levels of reported resolution. We identified 925 bacterial ribosome structures with a sufficiently high reported resolution that had been deposited in the PDB along with the corresponding cryoEM density map deposited in the EMDB (22). Performing hierarchical resolution calculations on these structures provides a real-world demonstration of the quantification of biomolecular structural information content, using the bacterial ribosome as an experimental case study.

To avoid complexities of developing a cryoEM-specific implementation of hierarchical resolution (*e.g.*, element-specific kernel weights, effects from charged atoms, electron beam-induced damage), we performed an initial simplified, technique-agnostic, hierarchical resolution calculation (see *Materials and Methods*). This implementation utilized an isotropic forward-model where each atom had the same density kernel weight (*ω_i_* = 1). Overall, this isotropic forward model is consistent with commonly used assumptions in models of electron microscopy imaging physics (38–40). Moreover, it can also be usedfor other imaging techniques (*e.g.*, atomic-force microscopy) and even X-ray crystallography, if the phases are known. To avoid representing higher-tier structural elements with isotropic forward-models, only the *d = 0* and *d = 1* levels of the hierarchy were considered (Eqn. 8). Consequently, the analysis was limited to bacterial ribosome structures with reported resolutions less than 5 Å. Fine tuning this implementation to capture technique-specific details and use the full structural hierarchy is a promising direction for future studies.

The results of the simplified, technique-agnostic hierarchical resolution calculations on all of the high-resolution, cryoEM-derived, bacterial ribosome structures in the PDB align well with our scientific intuition about structural biology experiments of biomolecular complexes (Fig. 4). At the per-residue level, the probability distribution for the *m^th^* residue containing atomic information, *P_m_*, is largely bimodal (Fig. 4A). *P_m_* takes on values of either 1 or 0, corresponding to the presence or absence of atomic information in the cryoEM density map, respectively. Notably, atomic information is present for at least some residues at every reported resolution in the calculation range (1.5 to 5 Å), but even at the smallest values of reported resolution, there were still residues without atomic information. This could be due to experimental factors, such as electron beam-induced damage (*e.g.*, decarboxylation of aspartic or glutamic acid), conformational dynamics, complications from preferred-angle issues, heterogeneous post-translational or post-transcriptional modifications, or even mismodeled residues. The atomic information content of only a few residues was ambiguous (0 < *P_m_* < 1); notably, at ∼0.7 in the range of 3.5 to 5 Å. This is likely a statistical effect for regions with extremely low SNR (*c.f.,* Fig. 3C). Most importantly, the proportion of residues that contain atomic resolution information (*i.e.*, *P_m_* ≈ 1) increases as the value of reported resolution decreases. In general, high spatial frequency features may be present in the data, but it is not *a priori* clear how they relate to the local structure of the biomolecule. Thus, this observation is a quantitative demonstration that better reported resolution makes it easier to localize biomolecule-specific structural information within the data (see *Discussion*).

**Figure 4.**
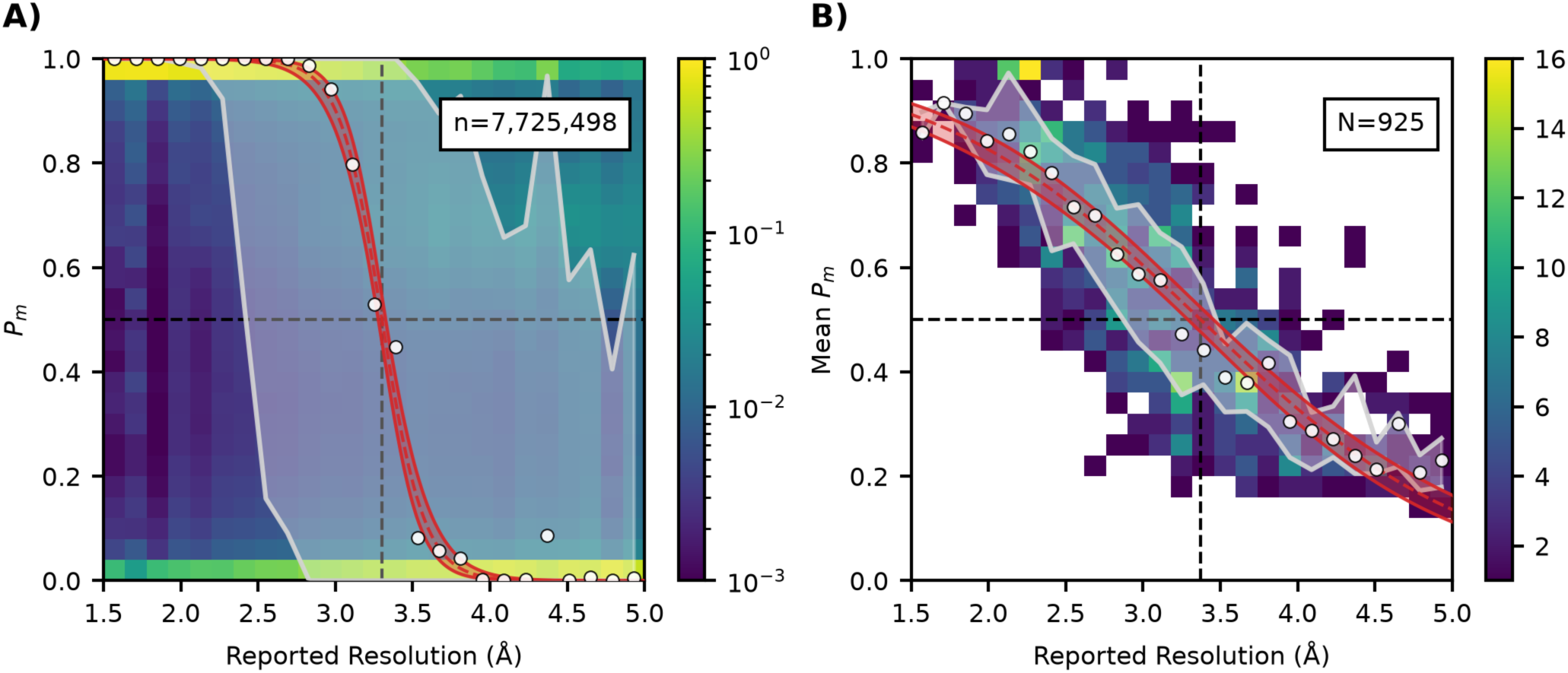
Hierarchical resolution *vs.* reported resolution for cryoEM-derived bacterial ribosome structures in the PDB. (A) Histogram of *P_m_* for all the 7,725,498 residues of 925 single-particle cryoEM-derived bacterial ribosome structures deposited in the PDB. The histogram maps the density of *P_m_* values normalized for each reported resolution bin. (B) Histogram of the mean value of *P_m_* calculated for each structure. The histogram maps the number of structures in each bin, and is not normalized. (A,B) The white regions indicate the interquartile range (25% to 75%), and the white dots indicate the median at each reported resolution. The median values were fit with a decreasing logistic function; red dashed lines show the best-fit solution, and the red region indicates the approximate 95% confidence band. Dashed black lines mark the best-fit logistic function midpoint at 3.3 Å (R^2^=0.99) and 3.4 Å (R^2^=0.97), respectively.

At the individual structure level, the mean value of all *P_m_* within each structure exhibits a unimodal monotonic relationship with reported resolution (Fig. 4B). Similar to individual *P_m_* values, the mean *P_m_* value increases with decreasing values of reported resolution. This demonstrates that increasing amounts of atomic information are embedded in entire experiments as the reported resolution improves. Surprisingly, even at the worst values of reported resolution near 5 Å, the bacterial ribosome structures deposited in the PDB contained approximately 20% of the possible amount of atomic information. That a significantly non-zero amount of atomic information is present at reported resolutions quite far from traditional notions of “atomic resolution” is not completely unexpected; altogether, this reinforces the view that atomistic models built into even “low resolution” cryoEM density maps can provide relatively “high resolution” structural information.

At both the residue and structure level, the median probability of the experimental data containing atomic information was well-fit by a logistic function (Fig. 4; red curves). The midpoints of the logistic functions were 3.31 Å and 3.37 Å, respectively. Thus, the experimental data contains atomic information for the majority of residues within a structure when the reported resolution is approximately 3.3 Å or better. However, for the experimental data to contain atomic information about all the residues in a biomolecular complex, the reported resolution must be even better—on average, only 90% of the residues will at 1.44 Å. This clarifies some ambiguity about casual uses of the terms “atomic resolution” and “near-atomic resolution” (17, 18). For atomic information about all of the residues, the reported resolution must be near 1 Å; however, for atomic information about the majority of the residues, the reported resolution must be near 3 Å (Fig. 4). Thus, the two terms are useful for discussing whether all of, or a majority of, the possible atomic information can be extracted from the data. Notably, for an extremely high-resolution X-ray crystallography-derived structure of rubredoxin at 0.43 Å (59), this simplified, isotropic hierarchical structure calculation revealed that every residue in that structure had *P_m_* = 1.0 (see *Supporting Material*). This demonstrates that complete atomic resolution is experimentally achievable. However, certain experimental situations, such as the presence of static molecular heterogeneity (*e.g.*, post-transcriptional or post-translational modifications) or dynamic molecular heterogeneity (*e.g.*, thermal motion), may make complete atomic resolution fundamentally impossible to achieve, especially if a structure is modeled with a single atomistic conformation for data that contains evidence for multiple conformations.

The logistic relationships between atomic information and reported resolution investigated here for cryoEM-determined bacterial ribosome structures demonstrates that hierarchical resolution calculations provide a representation of the structural information content of biomolecular structures as a distribution, which is richer than the single value of a reported resolution. A similar effect can be seen in the increased loss of information entropy with better reported resolution (Fig. S3), demonstrating the information theoretic basis of how hierarchical resolution calculations quantify the structural information content of an experiment (see *Supporting Material*).

## DISCUSSION

From the top down, the partition tree can be thought of as a mechanism to geometrically decompose the structure of a biomolecule into simple components with a significant positional locality (Fig. 1). From the bottom up, it defines the type of mutual, positional information that is shared by specific groupings of atoms. From both perspectives, the partition tree defines structural information through groupings of atoms in a manner that captures their intrinsic positional similarities. If a specific grouping is resolved, its positional information can be quantified by localizing the corresponding structural element. Thus, the concept of resolution is a question of which grouping along the hierarchy is most justifiably localized (see Eqn. 3).

As a concrete example of the positional localization of such a structural grouping, it is useful to consider single-molecule localization microscopy (SMLM). In SMLM, the detrimental effects of diffraction are bypassed by localizing a chromophore to a single point in space—the average location of its transition dipole moment. Thus, all the atoms in that chromophore form a single grouping whose positions are co-localized to that point. Furthermore, the atoms of any biomolecule(s) conjugated to that chromophore are also considered to be co-localized with this group. Thus, SMLM can be thought of as a reduced form of hierarchical resolution where a single HSR, representing the entire chromophore-conjugated biomolecule(s), is resolved by its localization to a single point in space. By itself, however, this single grouping does not provide much structural information other than the existence of the biomolecule(s). In a full treatment of hierarchical resolution, alternative groupings (*i.e.*, more granular HSRs) provide the increased-complexity representations necessary to resolve additional types of mutual positional information for the structural elements that comprise the biomolecule(s).

It is important to recognize that resolving biomolecular structural information in some experimental data is a fundamentally different problem than correcting that data for an instrument response function. If experimental data could be perfectly corrected to account for the distortions induced by the imaging processing (*e.g.*, by deconvolving with the true PSF), the question of which HSR is the best representation of the biomolecular complex would still remain unanswered. For example, even with perfectly corrected data, molecular heterogeneity caused by a thermal distribution of isoenergetic conformations (*e.g.*, of an intrinsically disordered protein) might obscure the definitive localizations of atomic positions without affecting the relative positional similarities captured by higher-level structural elements, such as domains.

This understanding of resolution also clarifies the ambiguity in the use of the terms atomic and near-atomic resolution (17, 18). The numbers typically used to represent “resolution” in techniques such as X-ray crystallography (nominal resolution, d_min_) or single-particle cryoEM (gold standard FSC resolution) are primarily quantifications of the instrument response (*e.g.*, the PSF and amount of noise) (60). These quantities are only correlated with atomic resolution in the sense that larger values make it more difficult to localize atomic positions. Distinct from this ease of localization, atomic resolution is a local property achieved when the individual positions of atoms in a residue are more justifiably localized than their common positional information as a composite residue (see Eqn. 8). In experimental situations described as “near-atomic resolution” (*e.g.*, nominal or FSC resolution ∼3 Å), atomic resolution can still be easily achieved (see Fig. 4) if the atomic-scale structural features of a residue generate a significant positional distinction amongst the atoms in the data. This explains the intuition behind why it should be easier to achieve atomic resolution for phenylalanine rather than alanine—the planar, electron rich phenylalanine sidechain is a positionally complex motif with more structural details that enable its localization in comparison to alanine with its simpler, methyl group sidechain. Resolution is, thus, an inherently local and compositional property of a biomolecule, and intuition about this aspect of its character is likely why the terms “atomic resolution” and “near-atomic resolution” are often used in a manner that conflicts with definitions of “atomic resolution” that are based solely on a cutoff to the nominal or FSC resolution.

The use of HSRs and structural hierarchies is not restricted to atomic or near-atomic resolution experiments. They can be employed to analyze nearly any type of experimental data that interrogates biomolecular structure. As such, it is worth asking: are there certain types of HSRs that are naturally suited to a particular technique? The most useful HSR for a particular technique depends upon how much structural information that technique can embed into the experimental data. For example, in a fluorescence resonance energy transfer (FRET) experiment, the FRET efficiency acts a one-dimensional proxy for the distance between a donor and acceptor fluorophore. Thus, the FRET efficiency can be used to localize a single degree of freedom (DoF). But how does this DoF map onto the hierarchical structure of the molecule? The answer requires an understanding of the structural DoFs of an HSR.

Partition trees for constructing HSRs treat biomolecular complexes as compressed entities comprised of multiple localizable structural elements that each have their own DoFs. The exact compression ratio depends on the HSR being considered. For example, for a prototypical bacterial protein consisting of ∼300 amino acids (∼9000 DoFs) organized into three domains, the purely tertiary level HSR, z = (*t11, t12, t13*), has nine DoFs—achieving an ∼1000x compression ratio. Six of these DoFs are inherently translational and rotational, while the remaining three DoFs correspond to the interdomain arrangement of the protein. Because each FRET signal can be used to localize a single DoF, a researcher could determine the geometry of the biomolecule using this HSR by performing at least three unique experiments (one per DoF). While a more detailed HSR would provide more structural information, the corresponding increase in DoFs would require significantly more experiments. Furthermore, the dynamic range of a FRET signal is typically effective for reporting on relatively large, nanometer-scale distances, such as the pairwise distances between tertiary structural elements (*e.g.*, *t_11_* to *t_12_*). FRET experiments are thus likely not sensitive enough to distinguish between, for example, different conformations of the constitutive residues of a loop. Thus, because of their low informational content and sensitivity, FRET experiments are ideally suited to inferring structural information using primarily quaternary and tertiary-level HSRs.

Similarly, SMLM experiments are best analyzed using quaternary-level HSRs; however, the use of antibodies to localize more detailed structural elements in conjunction with expansion microscopy is a promising future direction. Single-particle cryoEM experiments, however, are significantly more information-rich (Fig. S3), and are thus capable of handling atomically detailed HSRs. Hierarchical resolution is thus a very flexible framework that can be employed with a broad range of structural biology techniques to quantify biomolecular structural information content at the level naturally interrogated by a particular experimental technique. It is interesting to note that, while researchers can make decisions about which HSRs will best facilitate their analyses, similar considerations might apply to biomolecules engaging in biomolecular recognition (see *Supporting Material*).

Finally, the partition trees discussed above describe the geometries of static biomolecular complexes. From this perspective, the presence of any conformational dynamics during an experiment reduces the structural information content of the data and compromises resolution by inhibiting the localization of common positional information. Conformational dynamics can be accounted for, however, by defining multiple partition trees along a reaction coordinate. For example, in the case of a dynamic equilibrium between two metastable conformations, a unique partition tree can be constructed for each metastable state (Fig. 5). In such a case, both partition trees will originate from a common root node of the biomolecular complex itself (*i.e*., *C* in Eqn. 1), where the outermost bounding surface is large enough to encapsulate the biomolecular complex in either conformation. The result is a branched partition tree, and thus a branched PCFG. This branching does not need to occur at the root node—it can also occur at a lower level of *d* (*e.g.*, a loop with two conformations, or a residue with two rotamers; Fig. 5B).

**Figure 5.**
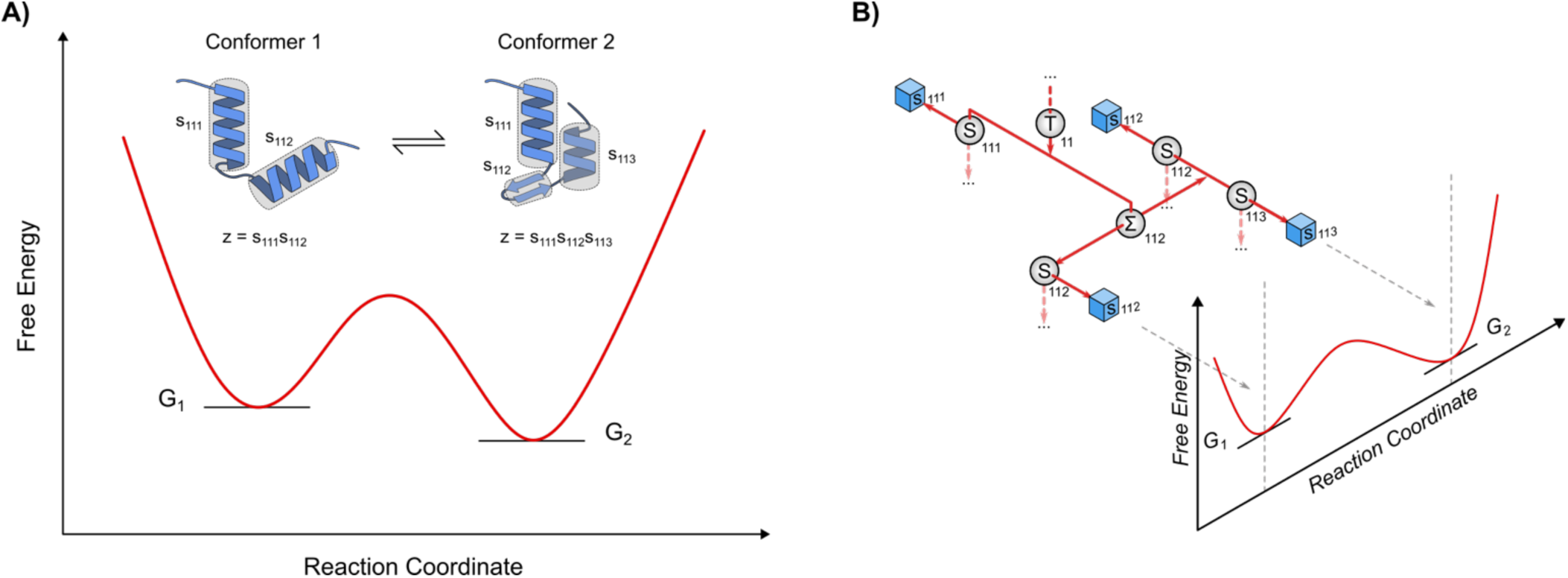
Effect of conformational dynamics on the structural hierarchy. (A) The free energy landscape for an example, two-state equilibrium between two biomolecule conformers. Partial HSRs describing the rearrangement of secondary structure elements are shown above. (B) Partial diagram of the PCFG showing the branching along the reaction coordinate of a virtual node, Σ_112_, into *S_112_* for conformer 1, or *S_112_S_113_* for conformer 2. The *S_111_* structural element is unaffected by the dynamics.

To systematize the grammatical organization of such branched PCFGs, we propose using non-terminal “virtual nodes” to represent the branched structural element independent of the instantaneous position of the molecule along the reaction coordinate for the motion of the corresponding structural element (Fig. 5B). While individual HSRs will contain the structural elements from only one of the branches emerging from the virtual node, the *E*_ℛ_ potential for the virtual node aggregates information from all its child branches. Thus, the virtual node can be used to better understand the resolution of dynamic structural elements within the structural hierarchy.

If a structural experiment enables the acquisition of unique datasets for each conformational state (*e.g.*, *via* classification during cryoEM density reconstruction), then each dataset can be analyzed using only HSRs from the branch at the appropriate point along the reaction coordinate. If the position along the reaction coordinate is *a priori* unclear, the posterior probability for HSRs from all the branches can be calculated using Eqn. 3, and the marginal probability for each branched structural element can be used to determine the most probable position along the reaction coordinate. If instead the data is ensemble and/or time averaged, then the molecule can be thought of as existing at multiple positions along the reaction coordinate. A terminal symbol for the virtual node may be added to the PCFG to represent this composite state. The forward model for such a virtual, composite structural element would be a mixture model (61), where the shape template (Eqn. 4) used to calculate the marginal likelihoods (Eqn. 3) is an ensemble and/or time weighted combination of the shape templates for the branched HSRs. Care must be taken, however, to ensure the mixture accurately reflects the thermodynamics and free energies along the reaction coordinate, for instance by using ensemble reweighting schemes (62).

## CONCLUSION

Here, we have developed a theory that connects biomolecular structural information at different lengths scales to resolution. This theory explains how a biomolecular complex can be decomposed using a partition tree to categorize the different types of structural information present on different length scales (Fig. 1). The resulting structural hierarchy of the partition tree implies a corresponding hierarchy of Rayleigh-like resolution criteria. Together, these two theoretical developments create a direct quantitative connection between structural knowledge about a biomolecule and biomolecular resolution.

The structural hierarchy demonstrates that biomolecular structural information can simultaneously exists on multiple length-scales, as long as they are resolved. This structural information can disappear, however, at points determined by the Rayleigh-like resolution criteria (Fig. 2). Importantly, these resolution criteria do not provide insight into when specific structural information appears or disappears, nor into which hierarchical level is the best representation of a particular biomolecular structure. Instead, they simply create observational bottlenecks at which points different emergent structural elements necessarily become the more relevant structural descriptions of a biomolecule.

Using the framework of the structural hierarchy and the corresponding set of Rayleigh-like resolution criteria, a researcher can determine the most relevant structural description of a biomolecule and quantify the structural information content of an experiment by performing a hierarchical resolution calculation (Eqns. 3-7). The fidelity of this calculation, how it quantitatively defines the structural information content of an experiment, and how it can be used as a local map-to-model quality metric for single-particle cryoEM was demonstrated using all the high-resolution, cryoEM-derived bacterial ribosome structures in the PDB (Figs. 4, S3). Hierarchical resolution itself is a detection sensitive resolution measure capable of quantifying local resolution (Fig. 3), and this clarifies the meaning of atomic resolution in structural biology experiments. Atomic resolution is an inherently local property of a residue; it describes when the positional distinction of the atomic-level structural elements in the data statistically outweighs their grouping within the higher-level structural elements that emerge along the hierarchy of biomolecular structure.

## DATA AVAILABILITY

Results and code to perform the two-dimensional image simulation analyses, and the bacterial ribosome hierarchical resolution calculations have been deposited in a Zenodo repository under the accession number 21536966 (20).

## AUTHOR CONTRIBUTIONS

C.D.K conceived the project, and C.D.K. and K.R. performed the research, analyzed the data, and wrote the manuscript.

## DECLARATION OF INTERESTS

The authors declare no completing interests.

## ACKNOWLEDGEMENTS

The authors thank Ruben Gonzalez, John Hunt, Ann McDermott, Eric Greene, Angelo Cacciuto, Riley Gentry, Erik Hartwick, Jason Hon, and Anjali Verma for helpful discussions. This work was supported by grants to C.D.K. from the National Science Foundation (CHE2442804, CHE2137630).

## SUPPORTING CITATIONS

References (63–67) appear in the *Supporting Material*.

## Notes

https://doi.org/10.5281/zenodo.10011335

### Competing Interest Statement

The authors have declared no competing interest.

### Summary of Updates

New section on applications to bacterial ribosome structures has been added; manuscript broadly updated; new supporting materials sections added.

