## Supporting Materials for "Biomolecular Resolution as a Quantification of Structural Information Content"

***TABLE OF CONTENTS***

1. Nearest neighbor residue calculations ………………………………………………………………... 2

2. Two-dimensional resolution measure sensitivity analysis ………………………………………………… 3

3. Hierarchical resolution calculations on X-ray crystallography-derived structures ……………………... 5

4. Information theoretic basis of biomolecular structural information content quantification …………….. 6

5. Applications of hierarchical resolution to biomolecular recognition ……………………………………... 8

6. References ……………………………………………………………………………………………….…. 9

### 1. Nearest neighbor residue calculations

An empirical reference distance to characterize secondary structure elements, $l_1$, can be made from the biomolecular complex structures deposited in the protein data bank (PDB). Specifically, the average distance between neighboring residues across all known structures is relatively agnostic of secondary structural context, and can thus provide a generalized reference distance. Using all the structures in the PDB, this distance can be calculated as the average distance from the center of mass (COM) of each residue to the COM of its closest neighboring residue. To ensure an accurate and a deterministic calculation, only structures with a reported resolution ≤ 4 Å, and that had been deposited before May 1, 2026 were included; additionally, the biomolecules in the structure had to be protein, DNA, and/or RNA polymers (*e.g.*, structures such as 1HYA or 4HP7 were excluded). Finally, only structures determined using cryogenic electron microscopy (cryoEM) or X-ray diffraction were included. These conditions yield N = 230,542 structures with a total of 240,818,990 residues. A histogram of the COM to nearest COM distances for those residues was constructed with 0.01 Å wide bins (Fig. S1). The average distance in this data is $\langle r \rangle$ = 4.33 Å, which is a reasonable choice for $l_1$ that yields $\sigma_1$ = 2.82 Å for isotropic Gaussian density kernels. Interestingly, the histogram exhibits structure with peaks and/or shoulders at distances of ~3.1, 3.5, 3.9, 4.1, 4.4, and 4.7 Å. Because of the order statistics involved in using the "closest" neighbor in this calculation (*i.e.*, minimum distance), each of these histogram features has a corresponding average value that is slightly longer. Presumably, these histogram features represent the

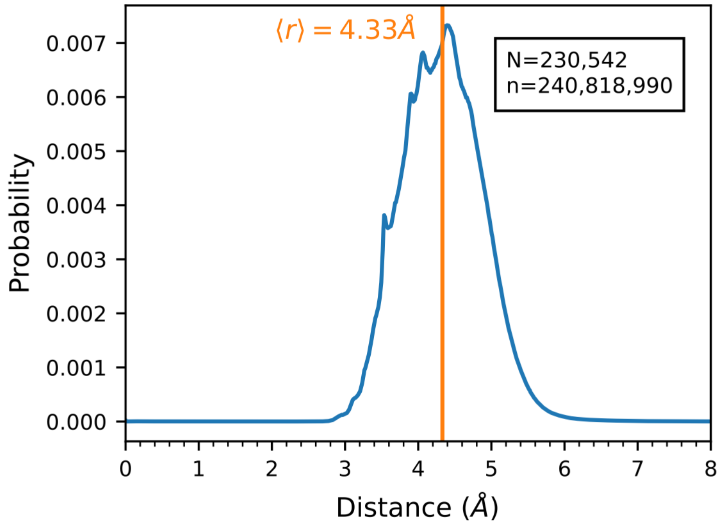


**Figure S1. Histogram of distances between nearest-neighbor residues.** CryoEM or X-ray diffraction derived structures deposited in the PDB with a reported resolution less than or equal to 4.0 Å were downloaded, and a histogram of the distances between the COM of each residue to the COM of its nearest neighbor was created.

unique secondary structure contexts found within the structures within the PDB (*e.g.*, alpha helix, beta sheet, b-form helix, *etc.*). Python code to perform this calculation has been deposited into a Zenodo repository (1).

### 2. Two-dimensional resolution measure sensitivity analysis

To better understand how the various parameters involved in a hierarchical resolution calculation affect the probabilities that are calculated, we performed additional two-dimensional simulation of the simplified molecule investigated in Fig. 3. In general, these simulations were performed as described in the *Materials and Methods*, but with a single parameter changed to investigate the effect of that parameter. The peak signal-to-noise ratio (PSNR) for the simulations in in Fig. S2A-E was 3, and in Fig. S2F was 10. For the anisotropic forward model discussed below, the density kernels were two-dimensional Gaussians with separate variances in the X and Y directions, and without a covariance term. Identical Rayleigh-like resolution criterion were used for the two directions, and marginalization of the widths for the two directions was performed using Gaussian quadrature. The additional marginalization dimension made simulations with anisotropic models significantly more computationally expensive. The number of simulated images *per* data point was 300 for Fig. S2A-E, and 100 for Fig. S2F.

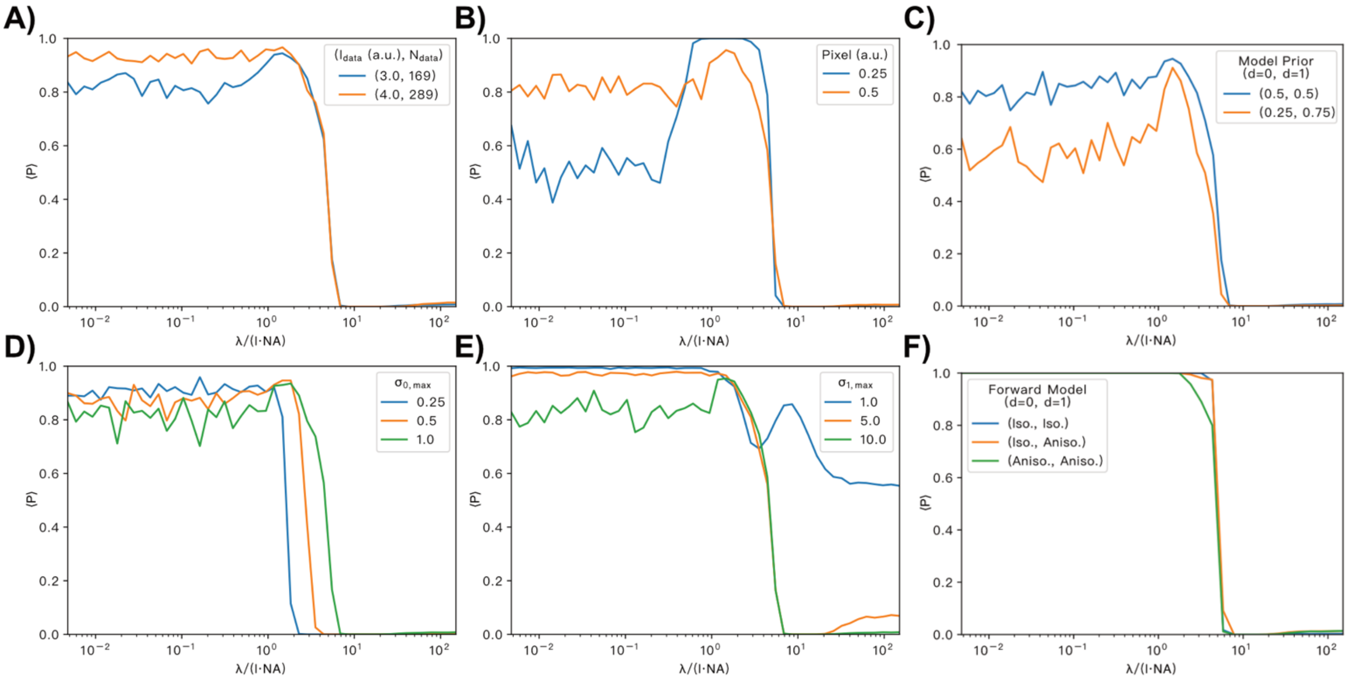


**Figure S2. Sensitivity analysis for two-dimensional simulations.** Plots of the probability that the simplistic, two-atom "molecule" is resolved at the atomic level as a function of normalized PSF width. (A) Effect of image dimensions and number of pixels. (B) Effect of pixel dimensions. (C) Effect of model prior. (D,E) Effect of integration bounds for Rayleigh-like resolution criteria: (D) $\sigma_{0,max}$, and (E) $\sigma_{1,max}$. (F) Effect of the use of isotropic and/or anisotropic forward models.

These additional simulations reveal several features of two-dimensional hierarchical resolution calculations. First, increasing the imaged region increased the number of data points in the inference calculation, and that increased the magnitude of the evidence for both models (Fig. S2A). Similarly, keeping the imaged region fixed but decreasing the pixel size increased the number of data points in the inference calculation, and that also increased the magnitude of the evidence for both model (Fig. S2B). It also shifted the Nyquist limit, which increased the evidence for the atomic model for normalized point-spread function (PSF) widths approximately down to the new Nyquist limit. Finally, as expected from the use of Bayes' rule for model selection (Eqn. 3), decreasing the model prior for the atomic model decreased the probability for the atomic model in a parameter-independent fashion (Fig. S2C).

Changing the limits of the Rayleigh-like resolution criterion for both the atomic and residue levels of the hierarchy had significant, although predictable, effects on the probability that the atoms were resolved at the atomic level (Fig. S2D-E). Increasing $\sigma_{0,max}$ primarily shifted the drop-off location of the binary resolution measure behavior towards larger values of the normalized PSF width (Fig. S2D). This demonstrates why the Rayleigh-like resolution criteria are so essential to hierarchical resolution. Without them (*i.e.*, with integration bounds for $\sigma_{0,max} = \infty$), there is always more evidence for the atomic-level model compared to any of the higher-level models. Thus, this demonstrates that a central feature of hierarchical resolution is the enforcement of the observational bottlenecks where other emergent structural elements necessarily become the more relevant structural descriptions of a biomolecule (see *Results*). Clearly, the exact definition of the Rayleigh-like resolution criteria is functionally important. Therefore, it must be justified from first principles as is done on the chemical level with the reference lengths and overlap criteria presented in the *Results* section (Fig. 2). On the other hand, increasing $\sigma_{1,max}$ increased the amount of evidence for resolution at the residue-level, but this was the highest level for this simplified two-dimensional molecule (Fig. S2E). As expected, when the residue-level structural elements were forced to be no larger than atomic-level structural element ($\sigma_{1,max} = \sigma_{0,max} = 1$), both the residue-level and atomic-level models were equally probable for normalized PSF widths so large that effectively no structural details exist. Notably, when the residue structural elements were allowed to be very large ($\sigma_{1,max} = 10$), additional evidence for the residue-level model was accumulated at smaller normalized PSF widths; most likely, in those cases, the two atomic puncta were interpreted as just being noise.

Finally, we investigated how the forward model affects the ability of hierarchical resolution calculations to quantify resolution (Fig. S2F). Given the apparent anisotropic elongation of an image of two atoms as they merge into a single density, we reasoned that an anisotropic forward model for the residue-level structural elements would equalize the improved performance of hierarchical resolution as a resolution measure over the Rayleigh or Abbe approaches (Fig. 3). Surprisingly, at least with high PSNR images, we saw no significant differences in the $\langle P \rangle$ *versus* normalized PSF width curves between the use of isotropic or anisotropic density kernels (Fig. S2F). One possible explanation is the "curse of dimensionality"—that the higher likelihood is balanced by the increased dimensionality of the parameter space (2). Regardless, these simulations demonstrate that the ability of hierarchical resolution calculations to quantify the structural information content of biomolecular complexes does not strongly depend upon the level of theory used for the imaging physics (*i.e.*, the choice of forward model). Rather, the use of hierarchical model selection to determine a spatial length scale inherently captures some emergent property of molecular structure—likely because those emergent structural details are intrinsic, local properties of the molecule, not the experiment.

**3. Hierarchical resolution calculations on X-ray crystallography-derived structures**

In X-ray diffraction experiments, the recorded data is the intensity of a diffraction spot, which is in reciprocal space. This contrasts with single-particle cryoEM experiments, where the recorded data is the cryoEM density (following reconstruction from the actually recorded individual micrographs), which is in real space. As discussed in the main text, hierarchical resolution calculations are performed in real space; so, hierarchical resolution calculations can be performed using cryoEM data, but cannot directly be performed using X-ray crystallography data. This issue is known as the phase problem of structural biology: atomistic model building for biomolecular complexes is performed using software that operates in real-space, but the data exist in reciprocal space and do not come with measured phases. Several well-established solutions to this problem exist. Phase determination is typically performed using complex auxiliary experiments (*e.g.*, anomalous scattering, isomorphous replacement), and/or *via* complex modeling approaches (*e.g.*, molecular replacement). In modern times, all X-ray crystallography-derived structures that are deposited in the PDB are required to include the structure factors (observed amplitudes), and the phases are explicitly included and/or can be calculated from the deposited atomistic model. Thus, both the amplitudes and phases of the structure factors are available for modern x-ray crystallography structures that are deposited in the PDB. Together the amplitudes and phases can be used to calculate electron density of the unit cell of the crystal. That electron density exists in real space. Thus, rather than having to develop a forward model for expressing a hierarchical structural representation in reciprocal space to perform a hierarchical resolution calculation, it is possible to compare the atomistic model with the computed electron density. With the simplistic, isotropic hierarchical resolution calculation developed in the main text, we performed such a calculation using an extremely high-resolution structure of rubredoxin ($d_{min}$ = 0.43 Å; PDB ID 30OR) (3). With this high level of reported resolution, along with the care taken by the authors to accurately produce an atomistic model from this data, we found that the probability of atomic resolution for every residue in this structure was $P_m = 1.0$, and thus that the information entropy *per* residue was H/N = 0.0 nats (see *Section 4*). Thus, applying hierarchical resolution calculations to such a high quality, X-ray crystallography-derived structure reveals that full atomic resolution is experimentally achievable, and that hierarchical resolution calculations can quantify that result.

**4. Information theoretic basis of biomolecular structural information content quantification**

In this section, information theory is used to explain the meaning of the "biomolecular information content" of a structural experiment. After explaining how a researcher's geometric understanding of a biomolecule's structure corresponds to an information entropy, we discuss how analyzing an experiment decreases that information entropy. The decreased amount corresponds to the biomolecular information content of the experiment. Finally, we demonstrate this concept using the high-resolution, single-particle cryoEM-derived structures of the bacterial ribosome analyzed in the main text.

For any particular biomolecule, there are many possible structural hierarchies, $\mathcal{H}$. All of those $\mathcal{H}$ are different probabilistic context-free grammars (PCFG), but they have something in common—they share the same partition tree, and thus have the same grammar. The $\mathcal{H}$ differ, however, by the set of the potentials for the PCFG rules, $E_{\mathcal{R}}$, which determine the probability of generating a particular hierarchical structural representation (HSR) using that $\mathcal{H}$. Thus, while the $\mathcal{H}$ share the same set of all possible HSRs, the probability of each HSR being the most likely structural description (given the researcher's previous knowledge and which experiment(s) are being considered) differs between the $\mathcal{H}$. For instance, before any experiments have been performed, the researcher might have no reason to favor one particular HSR over another. An initial structural hierarchy, $\mathcal{H}_i$, would therefore have a set of $E_{\mathcal{R}}$ potentials that yield equal model prior probabilities for each HSR. After having performed an experiment, a final structural hierarchy, $\mathcal{H}_f$, with $E_{\mathcal{R}}$ consistent with the model posterior probabilities from a hierarchical resolution calculation would favor certain HSRs (*e.g.*, those with atomic information). From this perspective, a structural biology experiment transforms the researcher's structural hierarchy from an $\mathcal{H}_i$ to $\mathcal{H}_f$. Thus, the information statistics of those two structural hierarchies quantifies the information gained by performing the experiment.

To be concrete, all the $\mathcal{H}$ considered in the simplified, isotropic hierarchical resolution calculations performed in the main text have $E_{Q_i \to q_i} = \infty$, $E_{T_{ij} \to t_{ij}} = \infty$, and $E_{S_{ijk} \to s_{ijk}} = \infty$. Within this background, a reasonable $\mathcal{H}_i$ that does not favor either the atomic-level or residue-level has $E_{R_m \to r_m} = E_{R_m \to a_m}$, such that $\beta \Delta E_m = 0$ for the *m^th^* residue. After having performed an experiment, a subsequent hierarchical resolution calculation like those discussed in the main text yields $P_m$ for each residue (Eqn. 8). With those results, a reasonable $\mathcal{H}_f$ is specified by equating the model posterior probability $P_m$ to the PCFG probability for the rule $R_m \to a_m$, which is $P(R_m \to a_m) = \left[1 + e^{\beta \Delta E_m}\right]^{-1}$, to yield the condition that $\beta \Delta E_m = \ln(1 - P_m) - \ln P_m$. Using these $\mathcal{H}_i$ and $\mathcal{H}_f$ structural hierarchies, the change in information entropy upon performing the experiment is $\Delta H = H_f - H_i$, where $H_f$ is the information entropy for $\mathcal{H}_f$ as calculated using Eqn. 2, and $H_i$ is that for $\mathcal{H}_i$. This expression is simplified by noting that $\beta \Delta E_m = 0$ for the initial structural hierarchy, and therefore that $H_i = N \ln 2$, where N is the number of residues in the structure. For the final structural hierarchy, the value of joint product $\beta \Delta E_m$ is determined by the value of $P_m$ alone, and there is no need to independently specify $\beta$ in Eqn. 2. Thus, for the types of hierarchical resolution calculations in the main text, the change in information entropy for the transition between $\mathcal{H}_i$ to $\mathcal{H}_f$ is

$$\Delta H = N \left\langle \beta \Delta E_m \cdot \left(e^{\beta \Delta E_m} + 1\right)^{-1} + \ln\left(1 + e^{-\beta \Delta E_m}\right) \right\rangle - N \ln 2$$

$$= -N \langle (1 - P_m) \cdot \ln(1 - P_m) + P_m \cdot \ln P_m \rangle - N \ln 2$$

where the angle brackets denote the average of the residues indexed by *m*. Since $H_f$ has a maximum value of $N \ln 2$ when all $P_m = 1/2$, the change in entropy is bounded between $-N \ln 2 \leq \Delta H \leq 0$. Thus, hierarchical resolution calculations result in a loss of information entropy, or rather a gain of certainty for the researcher's structural understanding of the biomolecule of interest.

We used the values of $P_m$ from the simplified, isotropic hierarchical resolution calculations performed on 925 high-resolution, single-particle cryoEM structures of the bacterial ribosome (Fig. 4) to calculate the loss of information entropy for each experiment (Fig. S3). The magnitude of this loss defines the biomolecular information content of the experiment. That the values of $\Delta H/\mathrm{N}$ were generally between approximately –0.5 nats to –0.693 nats, the latter of which is the greatest possible *per* residue loss, demonstrates that these experiments were all quite definitive in the (atomic- or residue-level) structural insight they provided into the bacterial ribosome. The median value of $\Delta H/N$ as a function of the reported resolution of the experiment exhibited a logistic relationship with a midpoint at 2.89 Å and $R^2$=0.89 (Fig. S3). Thus, while a significant amount of structural information is embedded into the data at poor reported resolutions above ~3.5 Å, the information content tended to increase as reported resolution dropped below 3.5 Å. Interestingly, this drop corresponds only to more information content; that content could be either atomic-level or residue-level information. From the plot of $P_m$ *versus* reported resolution, however, it is clear that the additional information content is primarily atomic-level information (Fig. 4). Regardless, there is only a small amount of information content gained from achieving reported resolutions less than ~1.8 Å (Fig. S3)—suggesting that factors other than data quality limit the interpretation of structural biology experiments of the bacterial ribosome (*e.g.*, conformational dynamics, model building issues,

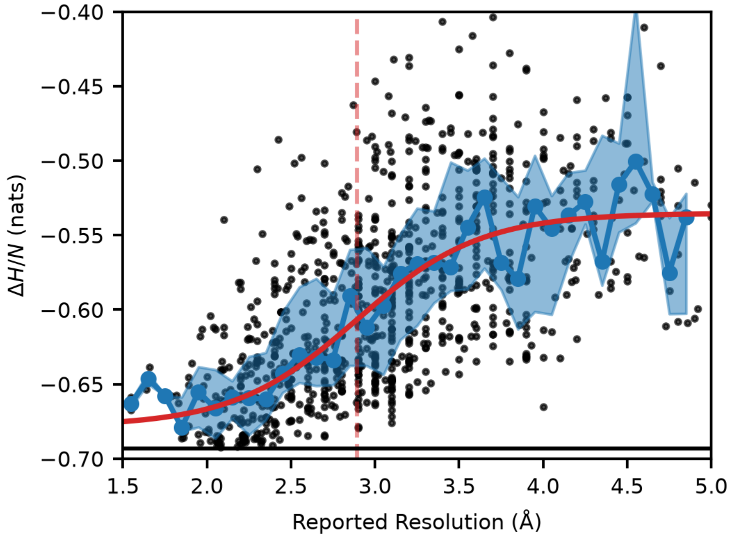


**Figure S3. Loss of information entropy for hierarchical resolution calculations of bacterial ribosomes.** The information entropy from the simplified hierarchical resolution calculations of the 925 high-resolution bacterial ribosome structures minus that for an initial PCFG in which both the atomic-level and residue-level representations are equally probable is plotted as a function of the reported resolution for each structure, normalized *per* residue (black dots). The median values within 0.1 Å-wide bins (blue line) are shown within the interquartile range for that bin (blue region). The best-fit logistic function (red line) to the median values had a midpoint of 2.89 Å ($R^2$ = 0.89).

electron beam-induced damage).

**5. Applications of hierarchical resolution to biomolecular recognition**

Since a hierarchical structural representation (HSR) is a description of particular structural information, HSRs can be used to identify and distinguish between different biomolecular complexes. Such a determination can be done by traversing the partition tree and comparing the HSRs for different complexes. For example, consider attempting to determine if an unknown complex, *X,* is a target trimeric complex, *Y*. If the highest-level HSR of *X* is $z = (q_1, q_2)$, then *X* cannot be *Y,* because the highest-level HSR of *Y* has three structural elements, $z = (q_1, q_2, q_3)$, not two.

Given the efficiency of such a search process, it is tempting to anthropomorphize biomolecular complexes and wonder if comparisons of different HSRs might also be used in nature for biomolecular recognition. If so, the recognized complex's partition tree would act to define the memory architecture for the recognizing complex, which would act as a biomolecular Maxwell's demon (4). Mechanistically, this requires the genes of the recognized and recognizing complexes to have co-evolved together. Furthermore, for such a mechanism to be possible, a slow internal degree of freedom (DoF) of the recognizing complex (*e.g.*, cis-trans isomerization of a proline) would need to act as a memory bit. The bit would store the binary identification status of a structural element of the recognized complex. For example, one DoF could encode the positive identification of the structural element for a domain. Interestingly, without any associated proofreading mechanism (5, 6), this mechanism places a thermodynamic limitation upon the accuracy of this recognition event: the stability of the internal DoF acting as a memory bit.

Higher fidelity could be achieved by simultaneously recognizing additional structural elements, and storing those identifications in additional internal DoFs. For example, the large dipole moment of an alpha helix could promote a conformational change to the DoF that successfully encodes the recognition of that helix. The recognition of more structural elements decreases the false-positive probability of accidently recognizing other structurally similar complexes in the cellular milieu.

Using this framework, molecular recognition events are equivalent to computations. Landauer and Bennet pointed out such calculations can be performed in a thermodynamically reversible manner (*e.g.*, first binding, followed by slowly writing the information to the DoF, and finally dissociation), whereas the erasure of the information does require energy dissipation (7). There are many molecular sources of free energy to use for that energy dissipation (*e.g.*, electrochemical gradients). Estimates from the use of hydrolysis of a single adenosine triphosphate molecule places an upper-limit of ~30 bits of HSR information that can be stored within a molecular complex (4); most likely the number is much lower. Regardless of how this dissipation is achieved, such a molecular memory would not be perfect. The stored information would thermalize over time, because the biomolecular complex dynamics are slaved to solvent modes (8). Additionally, the internal DoFs that serve as memory bits are anharmonically coupled to other DoFs in the complex (9), so stored information will slowly decohere. Finally, it is worth noting that any implementation of this framework in nature would likely use a different structural hierarchy than humans use to describe biomolecular structure.
